# SAMBA: A Segment Anything Model-based tool for semi-automated Behavioural Analysis of *Drosophila* and other model organisms

**DOI:** 10.1101/2025.08.21.671669

**Authors:** Sarah Mele, Long Nguyen, Joshua Millward, Jemma Gasperoni, Sebastian Dworkin, Zhen He, Travis K. Johnson

**Affiliations:** Department of Biochemistry and Chemistry, and La Trobe Institute for Molecular Sciences, La Trobe University, Bundoora, VIC, Australia, 3086; School of Computing, Engineering and Mathematical Sciences, La Trobe University, Bundoora, VIC, Australia, 3086; Department of Microbiology, Anatomy, Physiology and Pharmacology, and La Trobe Institute for Molecular Sciences, La Trobe University, Bundoora, VIC, Australia, 3086

**Keywords:** Segmentation, tracking, SAM2 model, behaviour, *Drosophila melanogaster*, zebrafish, *Danio rerio*

## Abstract

Quantitative behavioural analysis is a powerful approach for linking genotype to phenotype, but many existing tools require specialised hardware, extensive preprocessing, or coding expertise. We present SAMBA (Segment Anything Model for Behavioural Analysis), an open-access, Google Colab-based pipeline that harnesses the Segment Anything Model 2 (SAM2) for accurate, semi-automated tracking without thresholding or background subtraction. With minimal user input, SAMBA extracts movement parameters, detects behavioural states, and supports batch processing. Validating SAMBA in a *Drosophila melanogaster* model of 3-hydroxyisobutyryl-CoA hydrolase (HIBCH) deficiency revealed impaired locomotion, reduced speed, and altered decision-making, highlighting its ability to capture nuanced phenotypes in neurometabolic disease. We further demonstrate adaptability to adult *Drosophila* and larval zebrafish, underscoring its cross-species utility. By combining foundation-model segmentation with an accessible interface, SAMBA lowers technical barriers to high-throughput behavioural phenotyping and is readily extendable to diverse model organisms, life stages, and experimental paradigms. This flexibility positions SAMBA as a valuable platform for accelerating disease mechanism studies, genetic screens, and preclinical testing.

**SUMMARY STATEMENT:** We present an easy-to-use, open-access tool designed for *Drosophila* larval movement analysis, and readily customised for other applications, enabling rapid, scalable assessment of organism behaviour without specialised equipment or coding expertise.

## 1. INTRODUCTION

The fruit fly *Drosophila melanogaster* is a powerful genetic model organism that has contributed significantly to our understanding of neurobiology and diseases that impact the nervous system (Bellen et al., 2010; Jennings, 2011). A hallmark of many neurodegenerative disorders is progressive motor impairment, and the movement patterns of *Drosophila* larvae with equivalent genetic lesions reflect this. Larval locomotion normally follows predictable, stereotyped patterns, making deviations such as indecision or frequent changes in direction readily detectable (Jakubowski et al., 2012; Otto et al., 2018). Methods for examining larval locomotion patterns are particularly valuable in cases where diseases induce early lethality and preclude adult analysis. Assessing larval locomotion ability is also relevant for drug and toxicology experiments to determine adverse effects on neurological function (Li et al., 2019; Yap et al., 2021).

Existing methods for analysing larval locomotion often require specific hardware, extensive preprocessing, and substantial user input. Fiji/ImageJ plugins, centroid-based trackers, and threshold-dependent tools can be time-consuming, prone to segmentation errors, and sensitive to lighting or background variation. These limitations restrict throughput and make it challenging to capture subtle behavioural changes.

Recent advances in deep learning have transformed image segmentation across diverse domains including medical imaging (Chan et al., 2021), autonomous driving (Gupta et al., 2021) and agriculture (Logeshwaran et al., 2024), by reducing the burden of existing manual processes. However, applying these traditional segmentation models typically requires large, annotated datasets trained for a specific task, which can make them time-consuming and labour-intensive to develop. Foundation models such as Meta’s Segment Anything Model 2 (SAM2) overcome these constraints by being pre-trained on vast, heterogeneous datasets, enabling generalisation across domains without retraining (Ravi et al., 2024). In particular, SAM2 can segment objects with minimal prompting, even in complex visual environments, making it well-suited to biological applications without extensive computer vision expertise.

Here, we present SAMBA (Segment Anything Model for Behavioural Analysis), an open-access, semi-automated pipeline that uses SAM2 to track and quantify movement. It functions directly on raw video files without the need for thresholding, binarisation, or background subtraction - pre-processing steps common to other trackers. SAMBA is implemented in Google Colab, requiring no installation or coding skills, and supports batch processing for high-throughput studies. It extracts spatial and kinematic parameters, detects behavioural states such as pausing and head-casting, and outputs publication-ready visualisations alongside raw data.

To illustrate its utility, we apply SAMBA to a *Drosophila* model of 3-hydroxyisobutyryl-CoA hydrolase (HIBCH) deficiency, a rare neurometabolic disorder in humans (Loupatty et al., 2007; Taura et al., 2023). We demonstrate that SAMBA captures both gross locomotor deficits and altered decision-making behaviours in mutant larvae, providing phenotypic resolution beyond distance and speed metrics. We also show its adaptability to adult *Drosophila* and larval zebrafish, underscoring cross-species potential for behavioural phenotyping in genetic, pharmacological, and environmental studies. By combining the generalisation power of a foundation model with a user-friendly interface, SAMBA lowers technical barriers to quantitative behavioural analysis. Its flexibility makes it a valuable resource for researchers studying disease mechanisms, screening candidate therapies, and characterising behavioural phenotypes across a broad range of model organisms.

## 2. RESULTS

### 2.1 DEVELOPMENT of SAMBA for semi-automated larval tracking

We adapted the Segment Anything Model 2 (SAM2) to create SAMBA, a pipeline for high-accuracy larval locomotion analysis with minimal user input in the Google Colab environment. The notebook can be accessed via the GitHub repository (https://github.com/johnsonflygroup/SAMBA). To annotate an entire video, SAMBA requires a single frame to be annotated with point prompts for all objects of interest, and is capable of forward- and backward-time tracking. This allows rapid, flexible segmentation of larvae across diverse imaging conditions. We based its development on a set of 3-minute-long video recordings of 5 late 3^rd^ instar wild-type *Drosophila* larvae crawling in a circular arena of 60mm diameter (**Figure 1**). However, the approach of manually selecting each object of interest and feeding it into SAM2 should generalise to any animal tracking application without the need for training a dedicated object-specific detector. The tool includes an interactive calibration step to convert pixel distances into millimetres, ensuring outputs are directly interpretable in standard units provided that the camera is mounted perpendicular to the plane of object movement. From each segmented frame, SAMBA extracts position, orientation, and shape data, which are compiled into raw and summary CSV files along with annotated videos and track images (**Table 1**). We recommend running SAMBA on the Google Colab A100 or L4 GPU, capable of processing 3-minute videos (1080p, 30fps) tracking 5 objects in 20 and 30 minutes, respectively. Currently, the freely available T4 GPU option cannot process an equivalent video within the permitted 2-hour timeout period enforced by Google.

**Table 1.** Output files from SAMBA.

| Output folder | Output filenames | Description |
| --- | --- | --- |
| raw_frames | 0.csv...n.csv | Raw data for each frame including (x, y) coordinates, polygon size, ellipse major and minor axes, ellipse major/minor ratio, ellipse angle (degrees), major/minor ratio z-score, elongated/bent state, distance, and speed for each object. |
| problematic_frames | 0.csv...n.csv* | Frames copied from raw_frames which contain aberrant movement behaviour for each object. |
| good_frames | 0.csv...n.csv | Frames copied from raw_frames which contain no problematic frames (aberrant movement behaviour) for each object. |
| Paths | 0.csv...n.png | Movement tracings for each object. Includes highlighting for elongated and bent states. |
|  | output.csv | Aggregated data from good_frames. |
|  | tracking.mp4 | An annotated version of the original uploaded video with colour-coded, labelled polygonal overlays on each object. |

\*May not be present for each tracked object.

**Figure 1.**
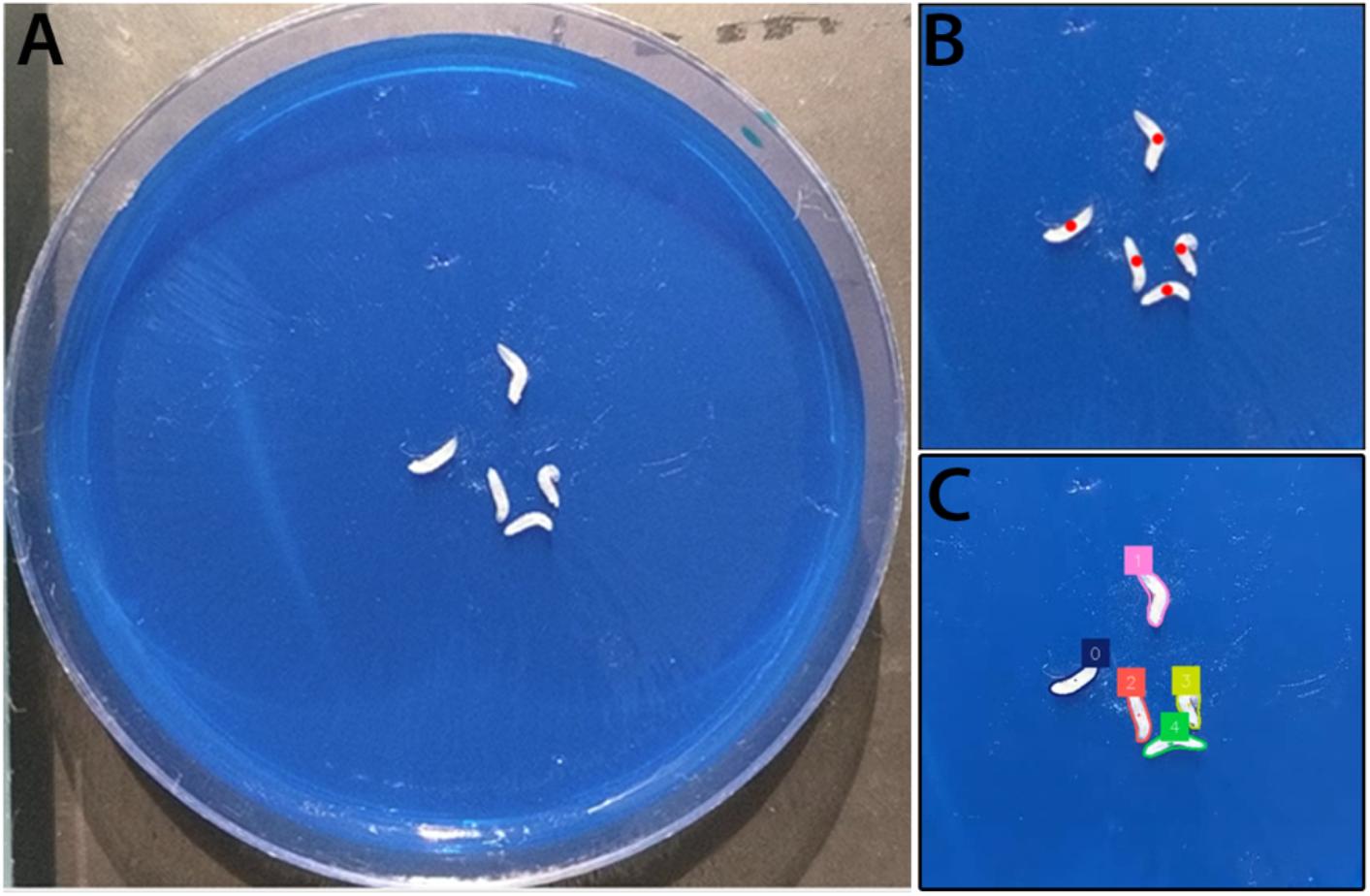
Manual larva selection via point prompting. (**A**) First frame of video of the larval arena used to apply SAM2 to larval locomotion. (**B**) User provided point prompts (red dots) to identify objects of interest. (**C**) Output masks from SAM2 with a colour-coded ID added to each larva.

*Drosophila* larval behaviours include *running*, which is a period of forward movement or uninterrupted crawling; *pausing*, when the larva ceases forward motion; and *head-casting*, where the larva sweeps its head side-to-side to sample its environment and make directional decisions (Berrigan and Pepin, 1995). We therefore enabled SAMBA to classify larval posture into ‘elongated’ or ‘bent’ states based on the ratio of ellipse major to minor axes (**Figure 2A**). These states correlate strongly with running versus head-casting behaviours, allowing posture-derived metrics such as speed and decision-making to be quantified (**Figure 2B-C**).

**Figure 2.**
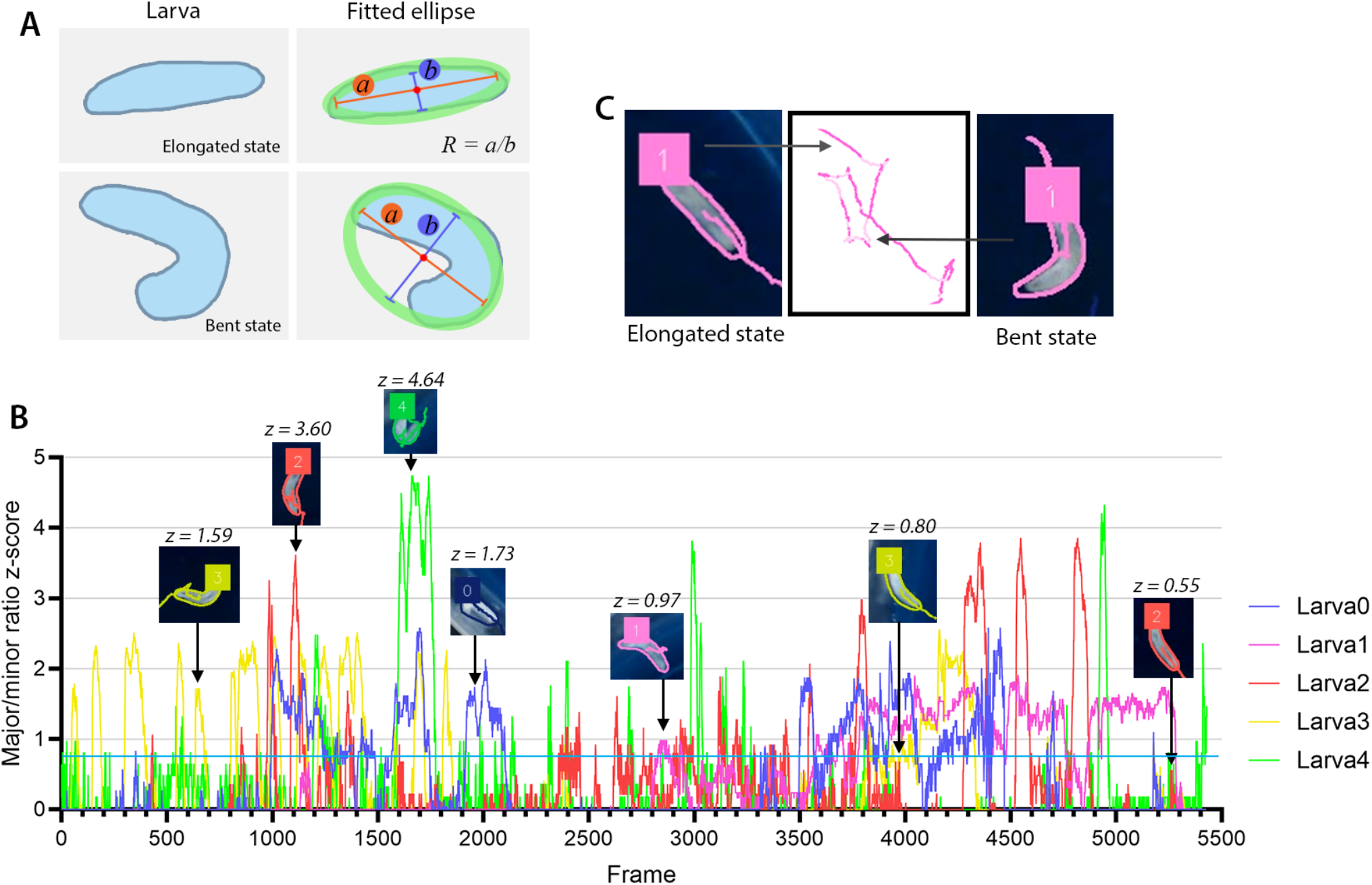
Characterisation of larval shape. (**A**) Schematic of how larvae are determined to be in a bent state using ellipses and the major-to-minor axis ratio (R = a/b, a = major axis, b = minor axis). (**B**) Plotted z-scores of R for 5 larvae. Examples of larva orientations at different z-scores are overlayed. The cutoff for the ‘bent’ shape was visually determined to be z ≥ 0.8. (**C**) Output images of larval tracks show darker coloured tracks when larva is elongated and a lighter colour when bent.

### 2.2 Quality control and batch processing

To minimise artefacts from segmentation errors, SAMBA flags specific larvae within each frame with outlier size (±4 standard deviations) or speed (±5 standard deviations) values and removes them from the aggregated model outputs (**Figure 3A-B**). The thresholds for outlier detection can be modified in an easily accessible code block in ‘1.2 Setup’. Note that these frames remain incorporated into the visual trace files. Cases of ID switching such as ‘merging’ (one object gradually merges into the ID of another object), ‘jumping’ (ID suddenly changes to another object), or ‘loss’ (an object gradually escapes its tracking polygon) are flagged by SAMBA, however user discretion should determine whether to omit the entire object from further analysis (**Figure 3C-E**). We further found that increasing the number of larvae per 60mm arena increased the frequency of ID errors (**Figure 3F**). Based on this, we recommend testing not more than 5 – 6 larvae in a single arena. For increased throughput, we included a batch processing mode for SAMBA, allowing users to process an entire directory of videos following point prompting in all videos prior to the initiation of tracking. Batch-processing file limits are constrained by the 2-hour maximum running time enforced by Google. An optimal run typically comprises five 3-minute videos tracking 5 larvae each, which comes to 1 hour and 40 minutes of total run time.

**Figure 3.**
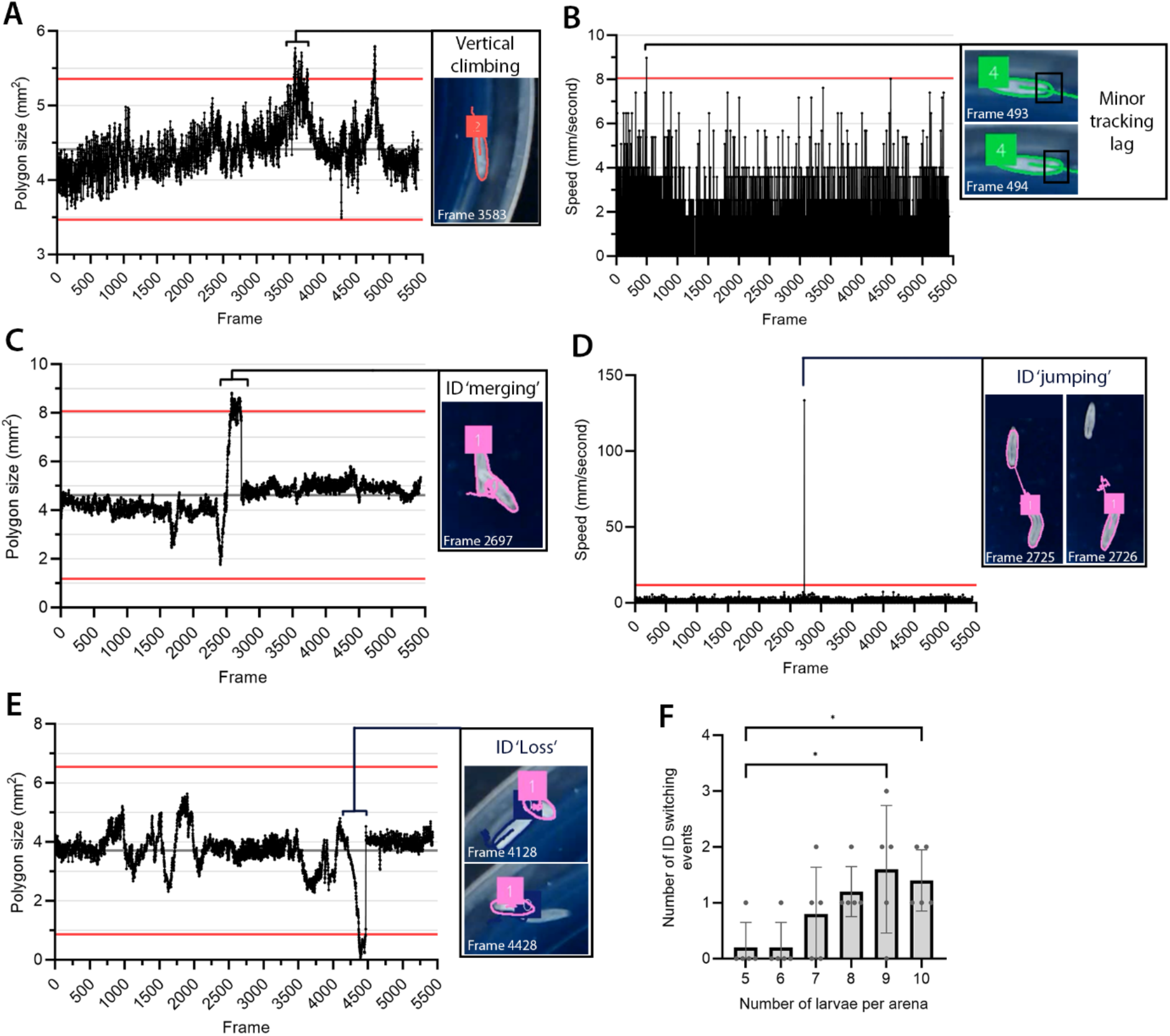
Quality control of SAMBA. Examples of tracking frames flagged as ‘problematic’ by SAMBA based on outlier size (**A, C, E**) or speed (**B, D**). Red lines indicate thresholds (size: ±4 standard deviations; speed: ±5 standard deviations). Sources of tracking errors include crawling vertically (**A**), tracking lag catch-up (**B**), ID merging (from collision, **C**), ID jumping (sudden high speed, **D**), or ID loss (gradual reduction in size, **E**). (**F**) Number of ID switching events (ID merging, jumping, loss) from 5 videos (3 minutes each) tracking up to 10 larvae in parallel. ANOVA *p < 0.05.

### 2.3 SAMBA for *Drosophila* disease modelling

To test the application of SAMBA in a *Drosophila* disease model, we selected a knockout allele of the gene encoding 3-hydroxyisobutyryl-CoA hydrolase (HIBCH), an enzyme required for the catabolism of the amino acid valine (Ferdinandusse et al., 2013). In humans, HIBCH deficiency is a rare autosomal recessive metabolic disorder resulting in mitochondrial abnormalities and subsequent neurological dysfunction (Soler-Alfonso et al., 2015; Taura et al., 2023) Flies mutant for the *Hibch* gene have recently been shown to have a strong locomotor deficit (Li et al., 2025). This together with our interest in *Hibch* made it a good candidate for demonstrating the functionality of SAMBA.

SAMBA identified strong locomotor impairment in *Hibch* mutant larvae in both overall distances travelled, and speed compared to age-matched controls (**Figure 4A-B**). SAMBA further identified that *Hibch* larvae spend longer in the ‘bent’ state compared to controls (**Figure 4C**), reflecting their short movement paths with frequent turns (**Figure 4D-E**), indicative of indecision or impaired exploratory control. After accounting for the time spent in the ‘bent’ state, *Hibch* larvae distance travelled and mean speed remained significantly reduced (**Figure 4G-H**). These findings are in line with adult locomotor deficits observed when *Hibch* is knocked down (Li et al., 2025). This analysis extends the known phenotypes for *Drosophila* HIBCH deficiency, highlighting the capacity of SAMBA to detect subtle, behaviourally relevant impairments.

**Figure 4.**
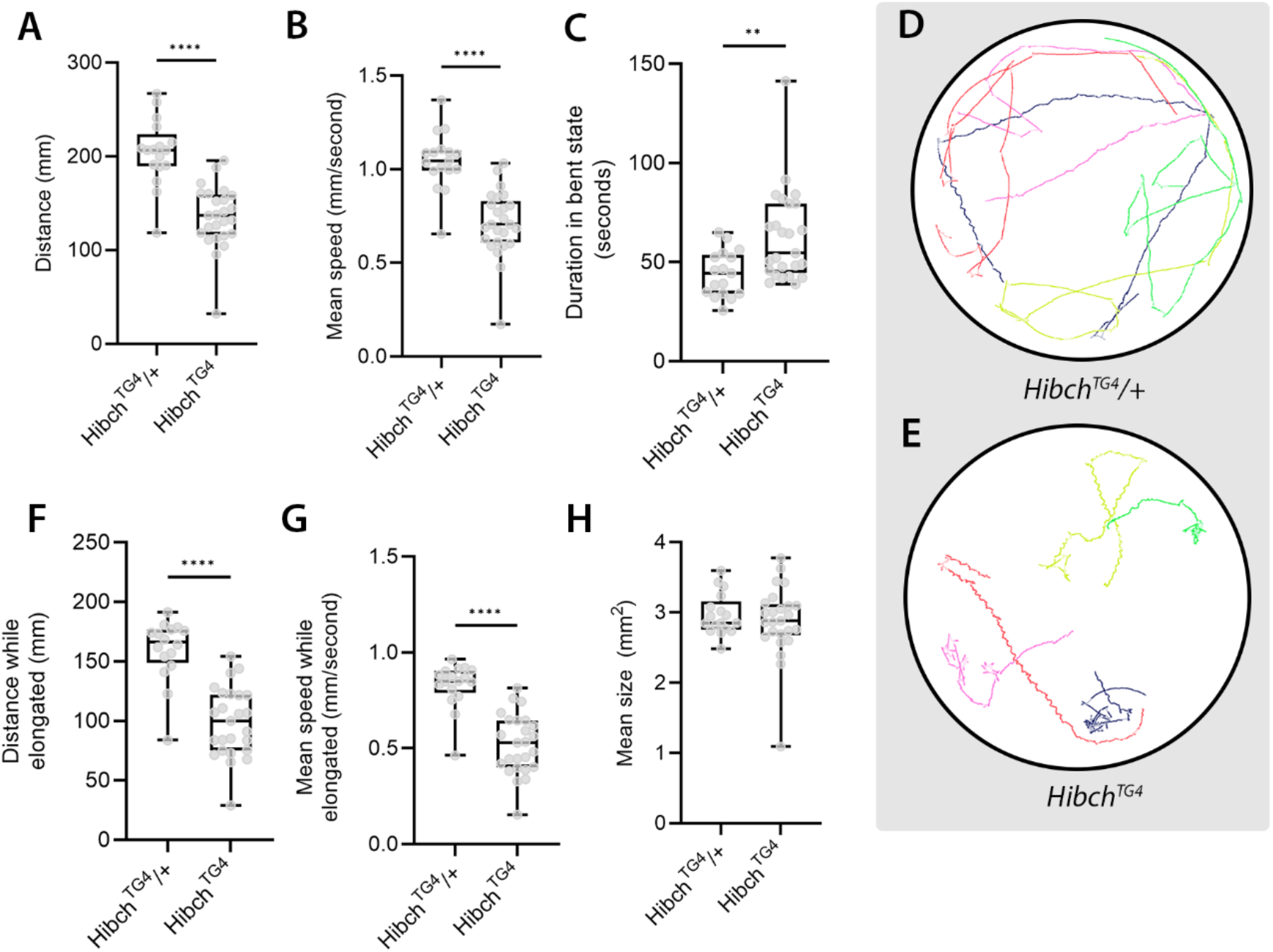
Locomotion deficit is observed in *Hibch*^*TG4*^ mutants. (**A**) Larvae homozygous for mutations in *Hibch* travel less distance and (**B**) are slower than heterozygous controls. Note these measurements depict overall larval movement regardless of orientation. (**C**) Larvae homozygous for mutations in *Hibch* spent more time stationary with increased head bending. (**D-E**) Traces of larval paths (5 larvae per plate), lighter line colour within individual paths indicate larva is in bent state. (**F**) Distance travelled and (**G**) mean speed adjusted for when larva is in elongated state only. (**H**) Size of larvae in locomotion analysis were not significantly different. Analysed using student’s T-test (****p < 0.0001, ** p < 0.01).

### 2.4 Adaptability of SAMBA to other models

Since SAMBA harnesses SAM2 for object tracking, it has potential for broad applications in behavioural analysis. As a preliminary test of this we trialled SAMBA for tracking adult *Drosophila* and zebrafish larval movement. Adult *Drosophila* are commonly analysed by a negative geotaxis assay, which is measured by tapping flies to the bottom of a vertical vial and recording the time taken for them to climb a defined distance up the wall, exploiting their innate escape response to gravity (Fergestad et al., 2006). SAMBA was capable of tracking individual flies and calculating their speed and distance travelled, supporting its potential for accurately quantifying this trait (**Figure 5A**). Finally, we also applied SAMBA to the analysis of zebrafish larvae (5 days post-fertilisation; dpf) housed individually in a 48-well plate, a widely used method for phenotyping genetic models, screening neuroactive compounds, or evaluating environmental or metabolic perturbations (Hernández et al., 2024; LaCoursiere et al., 2024; Lara and Vasconcelos, 2021). Here, SAMBA was able to track the larvae and capture speed and distance travelled in these fish (**Figure 5B**). These preliminary tests illustrate the flexibility of SAMBA for behavioural assays across model organisms.

**Figure 5.**
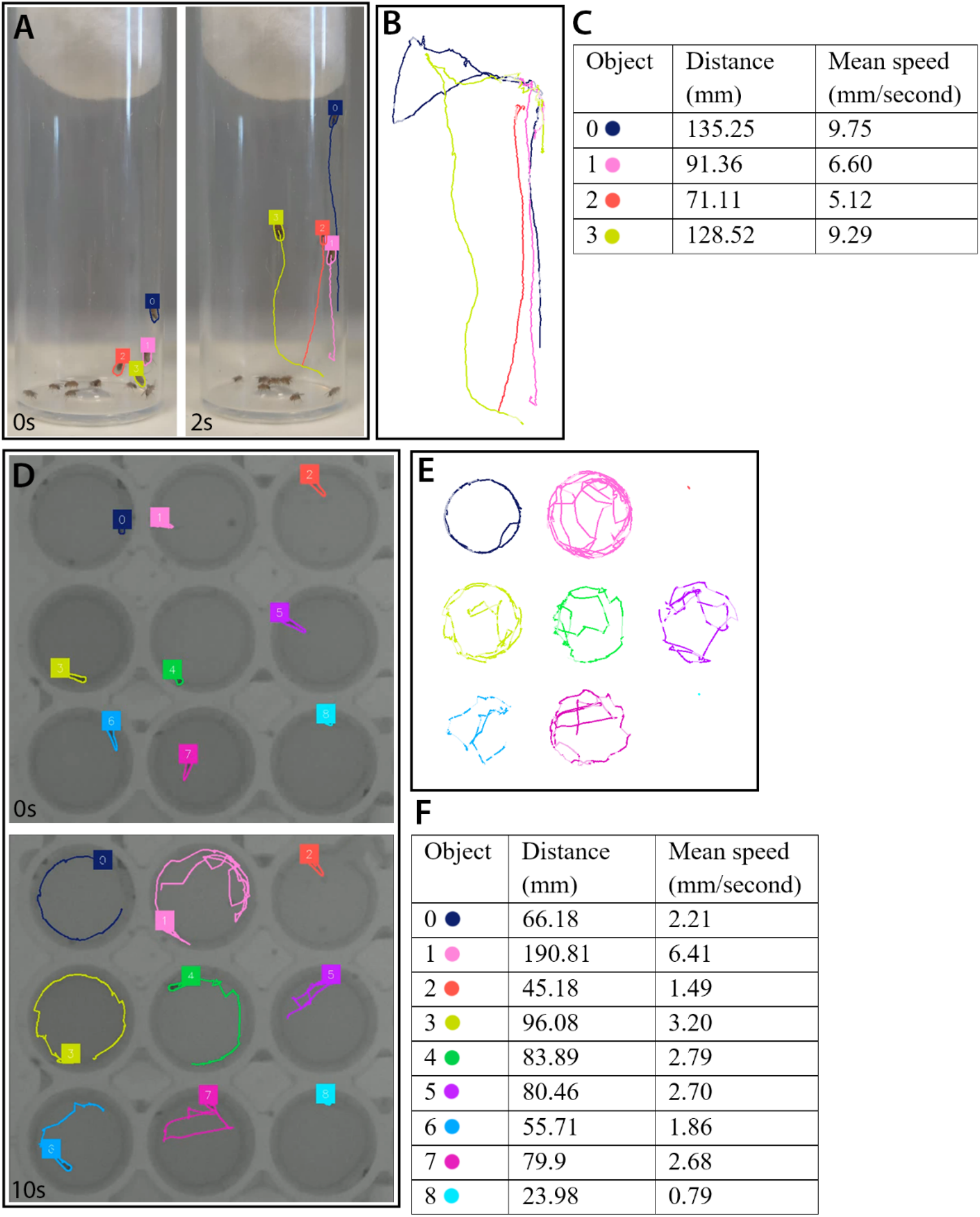
SAMBA tracking adult *Drosophila* and zebrafish larvae. (**A**) Example of adult *Drosophila* movement tracking. Flies were tapped to the bottom of the vial and the movements of a subset of flies were recorded for 14 seconds. Still images at 0 and 2 seconds (s) are shown with ID labels and movement paths overlaid. (**B**) Movement paths alone and (**C**) parameters of tracked flies. (**D**) Example of zebrafish larvae movement tracking. Zebrafish were recorded in 48-well plates for 30 seconds. (**E**) Movement paths and (**F**) parameters of tracked zebrafish.

## 4. DISCUSSION

Quantitative analysis of *Drosophila* larval locomotion has long provided valuable insights into neurological function, behaviour, and disease mechanisms. However, many commonly used approaches such as Fiji/ImageJ-based manual tracking, centroid-based tracking software (e.g. wrMTrck, Ctrax), or tools requiring binarised or thresholded input, can be time-consuming, require substantial user intervention, or depend on specific imaging conditions. While these methods have enabled detailed behavioural studies, they often involve trade-offs between throughput, accessibility, and behavioural resolution.

In this study, we introduce SAMBA, an open-access tool that harnesses the generalist capabilities of SAM2 to detect and track *Drosophila* larvae with no preprocessing and minimal technical setup. Our results demonstrate that SAMBA reliably identifies key features of larval movement, including continuous running, pauses, and head-casting behaviours which are components often associated with decision-making or sensory integration. Unlike many existing tools, SAMBA does not require thresholding or background subtraction to isolate animals, meaning raw videos taken directly from a smartphone, webcam, or other low-cost device can be analysed without preprocessing. The tool supports a range of common video file formats (including.*MP4*,.*MOV*, and.*AVI*), allowing for integration into diverse experimental setups. SAMBA features a batch-processing mode, allowing analysis of multiple videos concurrently, with no additional input once configured.

Our analysis of a *Drosophila* model of HIBCH deficiency illustrates SAMBA’s capacity to detect subtle locomotor phenotypes. We observed reduced distance travelled, slower average speed, and altered exploratory behaviours in mutant larvae, supporting its application for modelling complex neurometabolic conditions. This underscores its utility for detecting not only gross motor defects, but also more nuanced shifts in behavioural state and decision-making.

The flexibility and generalisability of SAM2 opens the possibility of adapting this pipeline to other model organisms (e.g. adult zebrafish, C. elegans, or even small vertebrates) as shown by our preliminary tests with adult *Drosophila* and zebrafish larvae. SAMBA can be adapted to different contexts using the modifiable parameters embedded in the script (ellipse R ratio cut-off, outlier detection). More specific modifications are possible for experienced coders due to directly accessible code. This makes SAMBA a versatile platform for applications in genetic screens, nutritional studies, pharmacological testing, or environmental stress experiments.

SAMBA runs via Google Colab, requiring no prior programming experience and only basic computer literacy, thereby making it easily accessible to general users. Users can upload their own videos and extract movement parameters through a guided interface. Unlike traditional local installations or commercial behavioural tracking systems, no installation or software license is required. A current limitation, however, is the computational cost associated with running foundation models like SAM2, which require access to an advanced GPU for efficient processing. While Colab offers a convenient hosted environment, users currently require paid access more powerful GPU runtimes. The minimum access price as of August 2025 is AUD$15.13, and the cost of analysing one 3-minute video tracking 5 objects is approximately AUD$0.29.

We reason that using SAMBA may substantially reduce hands-on analysis time, enabling researchers to reallocate effort toward experimental design, data interpretation, or follow-up assays. SAMBA therefore offers an efficient, accessible, and flexible approach for larval movement analysis, well-suited to both high-throughput screening and detailed behavioural phenotyping. By lowering technical barriers, this tool may help broaden the scope of behavioural genetics studies, particularly in models of neurological disease.

## 3. MATERIALS AND METHODS

### 3.1 *Drosophila* lines and maintenance

The following fly stocks were obtained from the Bloomington (BL) *Drosophila* Stock Center: *y*^1^, *w*^1118^ as the wildtype strain (BL6598); *y*^1^ *w*\*; *Hibch*^*TG4*^ /TM3,Sb^1^Ser^1^ (BL93364); and P150/ChFP-TM3, Sb (mCherry fluorescent protein (ChFP) balancer) (BL35524). The *Hibch*^*TG4*^ allele was balanced over TM3, ChFP and mutant larvae were selected based on the absence of ChFP using a Leica M165FC fluorescent stereo microscope. Flies were maintained on standard sugar-yeast food at 25°C in 12h:12h light:dark conditions.

### 3.2 *Drosophila* larva staging

Embryos were collected from population cages containing 80 females and 20 males and allowed to lay for 8h on apple juice agar supplemented with yeast paste. Eggs were collected and washed with distilled water and ∼10µl of embryos were deposited onto vials containing sugar yeast fly media incubated at 25°C in 12h:12h light:dark conditions until larvae reached the 3^rd^ instar wandering phase.

### 3.3 Zebrafish husbandry and embryo collection

Wild-type Danio rerio (Tübingen strain) were maintained under standard laboratory conditions at La Trobe University, in accordance with institutional animal ethics approvals (AEC16-091, AEC23-011). Adult zebrafish (>3 months old) were housed in a Tecniplast recirculating aquaculture system and maintained under a 14:10 hour light:dark cycle. To induce spawning, males and females were separated overnight using a transparent divider. Following a 10-hour dark period, lights were turned on to simulate dawn, and the divider was removed to allow mating. Fertilised embryos were collected within 30 minutes post-fertilisation, rinsed in E3 embryo medium (5 mM NaCl, 0.17 mM KCl, 0.33 mM CaCl_2_, 0.33 mM MgSO_4_), and incubated at 28°C.

### 3.4 Video acquisition

For *Drosophila*, five wandering 3^rd^ instar larvae were removed from the side of the vial using a blunt metal probe and transferred to the centre of a 60mm petri dish containing apple juice agar dyed dark blue (to enhance contrast). Larvae were left to acclimate to the dish for 3 minutes at room temperature. Five petri dishes per genotype were sufficient for this analysis. Petri dishes were positioned underneath a USB camera (Logitech HD Webcam) held by a retort stand at a height that captured the dish fully in the frame on a dark surface (see Lin et al. 2023 for setup schematic). A ruler was also captured in the frame for calibration. The lighting was adjusted using a lamp such that larvae were clearly distinguishable from the background. Three-minute videos were recorded. Statistical analyses of behavioural outputs was performed using GraphPad Prism (v10.4.2).

For adult *Drosophila*, 10 *w*^*1118*^ flies were placed into a wide vial and tapped to the base of the vial. Videos were recorded for 15 seconds on a Google Pixel 7a phone camera.

For Zebrafish, larvae were recorded at 5 dpf in 48-well plates, one larva per well in 1 mL E3 medium. Animals were acclimated for ≥30 min before testing. All experiments were performed in the DanioVision Observation Chamber (Noldus), with environmental temperature held at 28°C and data captured using a GigE digital camera with an infrared lens.

### 3.5 SAMBA development

SAMBA uses SAM2 to automatically generate a binary segmentation mask for each object in a frame. This mask is used to update the “inference state” of SAM2, which is used for generating predictions in subsequent frames. To extract object measurements, we first generated a polygon surrounding each object (using their binary masks), then computed the geometric centre (centroid) of each polygon (using the Python shapely library (Gillies et al., 2025). With the mask, polygon, and centroid, we were able to extract per-object measurements, such as the object pixel size (by summing up the number of foreground pixels in the mask), and various movement-based parameters, such as overall distance travelled, speed, and orientation (using the object centroid). These outputs were also used to generate different visualisations, such as the visual trace image of the object position across the entire video (using per-frame object centroids), and the video overlay outputs (using per-frame polygons and centroids).

To capture *Drosophila* larva decision-making, we used information on larva shape since head-casting behaviour involves body bending. After extracting object contours from the binary mask, we fitted an ellipse (OpenCV cv2.fitEllipse() function), which provided the ellipse centre point, major and minor axes lengths, and their orientation angles. The major-to-minor axis ratio (R) was used to determine if the larva was in an ‘elongated’ or ‘bent’ state. For classifying state in each frame, we applied a variable z-score threshold to the R distribution for each larva across the video. We found a threshold of 0.8 standard deviations was suitable to clearly distinguish the two states. We then integrated these states with movement data, allowing us to define speed, distance, and time spent in both ‘elongated’ and ‘bent’ states. Overlaying these on the trace images revealed excellent concordance between running behaviour and the ‘elongated’ state, as well as direction changes (decisions) and the ‘bent’ state.

By default, SAM2 holds the “inference state” for all video frames in memory, as it uses predictions from previous frames to inform predictions for the next frame. As such, this can lead SAM2 to exceed the available memory on a machine when there are many frames in a video. We particularly observed this with our larva videos, which were approximately 3 minutes long (approximately 5400 frames at 30 frames per second). To ensure SAM2 could process videos of any length, we reduced its memory usage by retaining only the most recent 1024 frames in the “inference state” (approximately 34 seconds of video at 30 frames per second). We observed that this provided a good trade-off between reducing memory requirements without causing noticeable deterioration in tracking outputs.

### 3.6 Running SAMBA

Instructions alongside screenshots are available in the supplemental files. Running SAMBA in Google Colab requires the use of a personal Google account, as outputs are stored in a Google Drive folder. In SAMBA, the required libraries and helper functions are initialised in Google Colab by running step 1 (Set-up environment), followed by the creation of working directories in Google Drive by running step 2 (Set-up working directory) which allows the notebook to read and write files to a personal Google Drive account. To determine the scale factor (mm/pixel), step 3 (Calibration) is run which prompts user selection of two points of known distance in the frame of a video. Alternatively, a predetermined scale factor can be used. The ‘Analysis Option A – Single video’ or ‘Analysis Option B – Batch processing’ sections are used to select a single video or a directory containing several videos respectively. In batch processing, each video is sequentially previewed and visible larvae on frame 0 (by default) are selected by clicking. Larva tracking was then completed automatically by SAMBA, and the outputs were generated at the specified directories. With minor changes and sufficient processing power, SAMBA could also be run locally.

## Acknowledgements

We thank the Bloomington *Drosophila* Stock Centre, and Professor Hugo Bellen and Dr Oguz Kanca (Baylor College of Medicine) for fly stocks, the Australian *Drosophila* Biomedical Research Facility (Ozdros) for stock importation, and members of the Johnson and He labs for helpful discussions.

## Data and resource availability

GitHub repository (https://github.com/johnsonflygroup/SAMBA).

## Author contributions

SM collected and analysed data, produced the figures and cowrote the manuscript. LN and JM wrote, edited and optimised the code and edited the manuscript. JG and SD performed the zebrafish work and edited the manuscript. ZH supervised the computational aspects of the work, contributed to algorithm optimisation, and acquired funding for the work. TKJ conceived the study, coordinated the project, supervised the biological components, acquired funding and cowrote the manuscript.

## Literature cited

Bellen, H. J., Tong, C. and Tsuda, H. (2010). 100 years of Drosophila research and its impact on vertebrate neuroscience: a history lesson for the future. Nat Rev Neurosci 11, 514–522.

Berrigan, D. and Pepin, D. J. (1995). How maggots move: Allometry and kinematics of crawling in larval Diptera. J Insect Physiol 41, 329–337.

Chan, H. P., Samala, R. K., Hadjiiski, L.M. and Zhou, C. (2020). Deep Learning in Medical Image Analysis. Adv Exp Med Biol. 1213, 3–21.

Ferdinandusse, S., Waterham, H. R., Heales, S. J. R., Brown, G. K., Hargreaves, I. P., Taanman, J.-W., Gunny, R., Abulhoul, L., Wanders, R. J. A., Clayton, P. T., et al. (2013). HIBCH mutations can cause Leigh-like disease with combined deficiency of multiple mitochondrial respiratory chain enzymes and pyruvate dehydrogenase. Orphanet J Rare Dis 8, 188.

Fergestad, T., Bostwick, B. and Ganetzky, B. (2006). Metabolic Disruption in Drosophila Bang-Sensitive Seizure Mutants. Genetics 173, 1357–1364.

Gillies, S., van der Wel, C., den Bossche, J., Taves, M.W., Arnott, J. and Ward, B. C., et al. (2023). Shapely.

Gupta, A., Anpalagan, A., Guan, L. and Khwaja, A. S. (2021). Deep learning for object detection and scene perception in self-driving cars: Survey, challenges, and open issues. Array 10, 100057.

Hernández, T. D. R., Gore, S. V., Kreiling, J. A. and Creton, R. (2024). Drug repurposing for neurodegenerative diseases using Zebrafish behavioral profiles. Biomed. Pharmacother. 171, 116096.

Jakubowski, B. R., Longoria, R. A. and Shubeita, G. T. (2012). A high throughput and sensitive method correlates neuronal disorder genotypes to Drosophila larvae crawling phenotypes. Fly 6, 303–308.

Jennings, B. H. (2011). Drosophila – a versatile model in biology & medicine. Mater Today 14, 190–195.

LaCoursiere, C. M., Ullmann, J. F. P., Koh, H. Y., Turner, L., Baker, C. M., Robens, B., Shao, W., Rotenberg, A., McGraw, C. M. and Poduri, A. H. (2024). Zebrafish models of candidate human epilepsy-associated genes provide evidence of hyperexcitability. iScience 27, 110172.

Lara, R. A. and Vasconcelos, R. O. (2021). Impact of noise on development, physiological stress and behavioural patterns in larval zebrafish. Sci. Rep. 11, 6615.

Li, J., Suda, K., Ueoka, I., Tanaka, R., Yoshida, H., Okada, Y., Okamoto, Y., Hiramatsu, Y., Takashima, H. and Yamaguchi, M. (2019). Neuron-specific knockdown of Drosophila HADHB induces a shortened lifespan, deficient locomotive ability, abnormal motor neuron terminal morphology and learning disability. Exp. Cell Res. 379, 150–158.

Li, Y., Wu, T., Li, Y., Xu, C., Zhou, C., Li, Z., Shang, W., Wang, L., Liu, Z., Wang, J., et al. (2025). Ectopic protein lysine methacrylation contributes to defects caused by loss of HIBCH or ECHS1. Cell Rep. 44, 115379.

Logeshwaran, J., Srivastava, D., Kumar, K. S., Rex, M. J., Al-Rasheed, A., Getahun, M. and Soufiene, B. O. (2024). Improving crop production using an agro-deep learning framework in precision agriculture. BMC Bioinformatics. 25, 341.

Loupatty, F. J., Clayton, P. T., Ruiter, J. P. N., Ofman, R., IJlst, L., Brown, G. K., Thorburn, D. R., Harris, R. A., Duran, M., DeSousa, C., et al. (2007). Mutations in the Gene Encoding 3-Hydroxyisobutyryl-CoA Hydrolase Results in Progressive Infantile Neurodegeneration. Am J Hum Genetics 80, 195–199.

Otto, N., Marelja, Z., Schoofs, A., Kranenburg, H., Bittern, J., Yildirim, K., Berh, D., Bethke, M., Thomas, S., Rode, S., et al. (2018). The sulfite oxidase Shopper controls neuronal activity by regulating glutamate homeostasis in Drosophila ensheathing glia. Nat Commun 9, 3514.

Ravi, N., Gabeur, V., Hu, Y.-T., Hu, R., Ryali, C., Ma, T., Khedr, H., Rädle, R., Rolland, C., Gustafson, L., et al. (2024). SAM 2: Segment Anything in Images and Videos. arXiv.

Soler-Alfonso, C., Enns, G. M., Koenig, M. K., Saavedra, H., Bonfante-Mejia, E. and Northrup, H. (2015). Identification of HIBCH Gene Mutations Causing Autosomal Recessive Leigh Syndrome: A Gene Involved in Valine Metabolism. Pediatr Neurol 52, 361–365.

Taura, Y., Tozawa, T., Isoda, K., Hirai, S., Chiyonobu, T., Yano, N., Hayashi, T., Yoshida, T. and Iehara, T. (2023). Leigh-like syndrome with progressive cerebellar atrophy caused by novel HIBCH variants. Hum. Genome Var. 10, 23.

Yap, Z. Y., Strucinska, K., Matsuzaki, S., Lee, S., Si, Y., Humphries, K., Tarnopolsky, M. A. and Yoon, W. H. (2021). A biallelic pathogenic variant in the OGDH gene results in a neurological disorder with features of a mitochondrial disease. J Inherit Metab Dis 44, 388–400.

